# Mechanistic interpretation of biological tissue growth experiments with a computational model

**DOI:** 10.64898/2026.03.12.711199

**Authors:** Shahak Kuba, Matthew J. Simpson, Pascal R. Buenzli

## Abstract

The growth rates of biological tissues are influenced by the existing substrate geometry, mechanobiological processes and the interplay between them. Disentangling the contributions of geometry and mechanobiology experimentally is challenging, as mechanical properties are difficult to measure and tissue samples provide only static snapshot in time. However, the composition of a tissue preserves cues of the dynamic processes that shaped its architecture. In this work, we present a computational model of tissue growth that captures aspects of the interplay between geometry, mechanics, and stochastic biological processes, which we use to generate synthetic tissue compositions directly comparable with experimental samples. This framework enables quantitative analysis of tissue morphology, inference of underlying growth mechanisms, estimation of dynamic rates from single-time-point data, and investigation of how stochasticity contributes to emergent growth patterns. We demonstrate the applicability of the model to simulate the growth of different tissue types by applying this framework to two distinct tissue growth scenarios: (i) tissue grown within 3D-printed porous scaffolds, and (ii) bone formation in cortical pores.

## 1 Introduction

The growth of biological tissues is central to the development, repair, and regeneration of organs and tissues. Tissue growth plays an important role for example in bone remodeling, tumor growth, and wound healing (Martin et al., 1998; Poujade et al., 2007; Rolli et al., 2012; Wozniak and El Haj, 2007; Lowengrub et al., 2009), as well as in tissue engineering and regenerative medicine, where biological tissue growth is involved in engineering the repair or replacement of damaged or diseased tissues (Hollister, 2005; O’Brien, 2011; Bidan et al., 2013). During tissue growth, new biological material is usually created within constrained spaces with experimental observations indicating that the geometry of both the constraining space and the existing tissue substrate influences the rate of growth (Rumpler et al., 2008; Bidan et al., 2012; Schamberger et al., 2023). The accumulation of new material within these geometric constraints induces crowding effects and mechanical stresses on individual cells, which may affect their ability to proliferate, differentiate, and survive through mechanobiological processes (Nelson et al., 2005; Discher et al., 2005; Lim et al., 2010; Ladoux and Mége, 2017). Additionally, cell-scale cues arising from the geometry of the existing substrate are also known to influence cell fate (McNamara et al., 2010; Dobbenga et al., 2016; Callens et al., 2020). The tissue growth dynamics arising from the interplay between geometry, mechanics, and cell fate in turn influence the rate of tissue growth, yet the precise quantitative details of these interplay remain poorly understood. Experimental approaches for studying the interactions between geometry and mechanics can often be challenging, especially as tissue samples provide single snapshots in time and offer limited insight into growth dynamics. However, the composition of a tissue contains useful cues that provide a record of some of the dynamic processes that led to the architecture of a tissue.

Biological tissues are primarily composed of an extracellular matrix (ECM) and differentiated cells embedded within it (Alberts et al., 2002; Kurniawan and Bouten, 2018). Since the material composition forms during growth, the properties of differentiated cells and ECM encode valuable information about the developmental history and can provide a fingerprint of the underlying processes that are regulated in healthy tissues, and dysregulated in diseased tissues. Confocal microscopy and quantitative backscattered electron imaging of tissue samples, such as those in Figure 1, provide a means to measure material properties, including spatial information relating to differentiated cells. Several studies use these measurements to characterize ECM properties such as fiber alignment as well as the shape, orientation, and number densities of differentiated cells within different tissue types (Roschger et al., 2008; Vatsa et al., 2008; Ascenzi et al., 2008; van Hove et al., 2009; Britz et al., 2012; Mader et al., 2013; Lanaro et al., 2021). Beyond providing a quantitative description of biological tissue composition, these data sets can be leveraged within mathematical modeling to help infer the mechanisms that govern tissue growth since these models can be designed to explicitly incorporate specific mechanisms. For example, mineral density distributions in trabecular bone have been used to study bone mineralization dynamics (Ruffoni et al., 2007), osteocyte density in cortical pores to estimate osteoblast burial rates (Buenzli, 2015), and mouse incisor enamel measurements to formulate predictive theories of cellular responses to strain variations (Cox et al., 2024). In this regard, mathematical modeling offers a powerful framework for generating and testing hypotheses, analyzing experimental data, and could be useful for disentangling interplay between geometry, mechanics, and cell fate in the processes underlying tissue growth.

**Figure 1.**
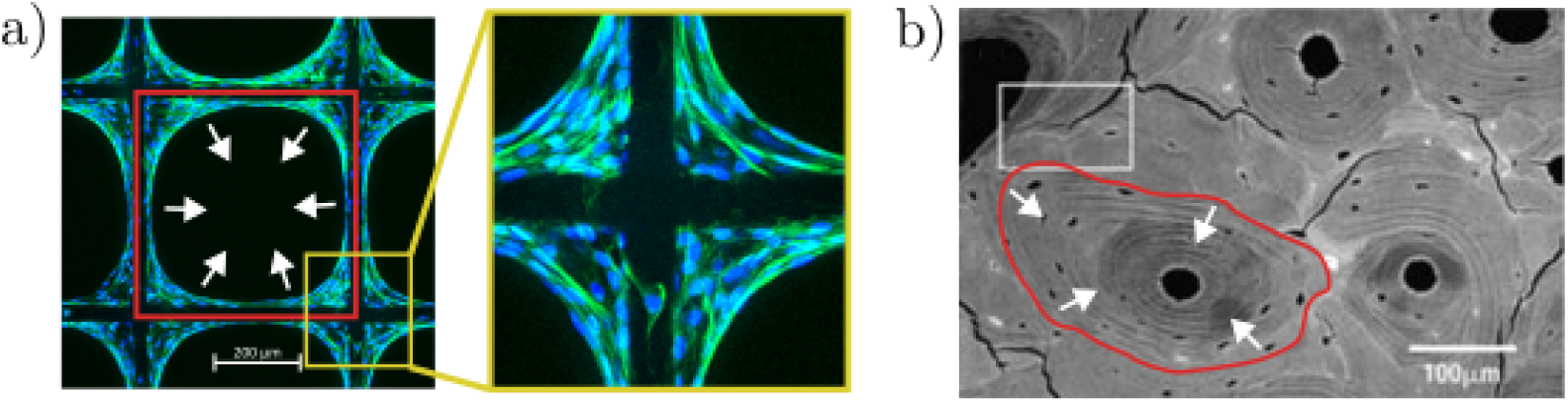
Images from case studies of biological tissue growth. (a) *In vitro* tissue growth of MC3T3 cells within a 400 μm × 400 μm 3D-printed square porous scaffold cultured over 14 days. The extracellular matrix (ECM) is depicted in green and differentiated cell nuclei are indicated in blue. (b) *In vivo* cross section of human bone tissue with multiples cortical pores highlighted in red surrounding the Haversian canal system with embedded cells (osteocytes) are indicated in black (reproduced with permission from Skedros et al. (2005)).

Several mathematical models of tissue growth have been developed to predict and help understand the evolution of biological tissues, with varying levels of biological detail. Phenomenological models capture the macroscopic behavior of growing tissues by defining growth laws that link the velocity of the tissue boundary and its curvature, for example, mean curvature flow (Rumpler et al., 2008; Guyot et al., 2014; Callens et al., 2023). Continuum mechanics approaches typically employ growth tensors to describe tissue deformation and expansion (Ambrosi and Guana, 2007; Dunlop et al., 2010; Goriely, 2007). However, these growth laws and growth tensors are difficult to relate to biological processes. Cell-based models, which include both continuum and discrete approaches, complement phenomenological descriptions by explicitly accounting for cell populations and modeling their associated behaviors (Bidan et al., 2012; Joly et al., 2013; Gahffarizadeh et al., 2018). These modeling frameworks have shown that some curvature effects arise from the crowding and spreading of cells (Alias and Buenzli, 2017; Hegarty-Cremer et al., 2021; Kuba et al., 2026). At the individual cell level, these models are also able to connect mechanical stresses to tissue organization and to cellular events (Baker et al., 2019; Murphy et al., 2019; Buenzli et al., 2025; Brown et al., 2025). While these modeling approaches have provided valuable insights into tissue growth, they tend to focus on the evolution of the tissue surface rather than generating a tissue composition that is comparable to experimental samples.

In this work, we present a computational model of biological tissue growth that can be used to simulate and analyze the dynamic processes that lead to observed tissue compositions in confined geometries. The model explicitly represents individual cells and includes mechanical interactions between cells, the generation of ECM, and incorporates cell proliferation, differentiation, and death as stochastic events. Unlike previous interface-based or curvature-controlled growth models, this framework explicitly records the positions and orientations of differentiated cells as tissue forms, enabling direct comparison of synthetic and experimental tissue compositions at single-cell resolution. Comparisons between model-generated predictions and experimental data enables us to: (i) relate mechanical interactions between cells to both their orientations and the shape of the tissue interface; (ii) identify key mechanisms that are important for regulating tissue growth; (iii) estimate cell differentiation rates at different times from single snapshot; and, (iv) analyze how modeled mechanisms and stochasticity shape tissue composition and give rise to random growth patterns. To demonstrate the flexibility of this framework, we apply it to investigate data from tissue engineering cell culture experiments in 3D-printed scaffolds, and bone formation in cortical pores.

## 2 Materials & Methods

This study is primarily simulation-based, using comparisons between computational model predictions and experimental observations to interpret the mechanisms underlying tissue growth. While experimental data provide quantitative description of tissue composition, they do not reveal the precise mechanisms that control the growth behaviour. By applying mechanistic simulation models to interpret these data, we obtain biological insights that are otherwise difficult to access.

### 2.1 Mathematical model

The mathematical model simulates tissue growth by evolving a discrete representation of the tissue interface, with newly formed tissue filling the region behind it. The interface is represented as a one-dimensional chain of *N* tissue-forming cells connected through their boundaries. The position of the *i*th cell at time *t* is given by the two-dimensional position vectors of its boundaries, ***r***_*i*−1_(*t*) and ***r***_*i*_ (*t*) (Figure 2a).

**Figure 2.**
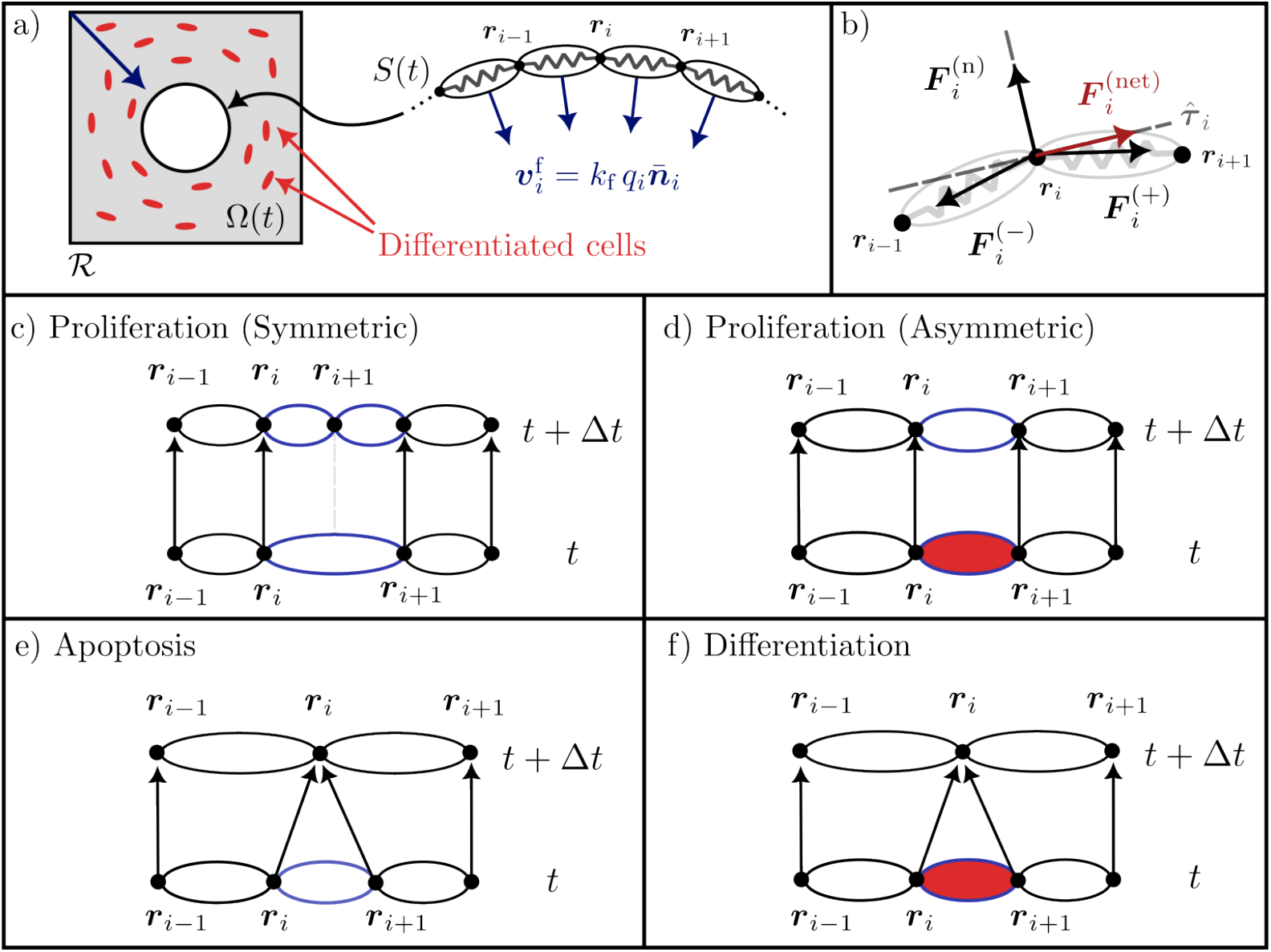
Overview of the mathematical model.(a) Tissue growth region *ℛ* and time-dependent tissue area *Ω*(*t*) showing differentiated cells (red) and the discrete cell-based tissue interface *S*(*t*). Tissue formation displaces interface cells perpendicularly in the direction 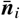 with a velocity proportional to the local cell density *q*_*i*_. (b) Mechanical force diagram at cell boundary ***r***_*i*_ showing two restoring forces from springs on either side and the normal reaction force as well as the net force. (c-f) Examples of discrete cells undergoing proliferation, asymmetric proliferation, apoptosis, and differentiation over a time step Δ*t*. Highlighted cells undergo the indicated process, and red cells denote differentiated cells as shown in (a).

Tissue growth arises from three processes: (i) mechanical interactions between neighbouring cells, (ii) ECM deposition at the interface, and (iii) stochastic cellular events. Following previous discrete cell–cell mechanical interaction modeling approaches, intercellular forces are represented by representing cell bodies as mechanical springs (Murray et al., 2009; Murphy et al., 2019; Buenzli et al., 2025). Cells are assumed to generate an area of tissue material at a constant rate *k*_f_ (μm^2^/day) that displaces them perpendicularly to the interface (Buenzli, 2015; Alias and Buenzli, 2017; Kuba et al., 2026). Together, these two mechanisms yield two independent velocity contributions that evolve the position of a given cell boundary ***r***_*i*_, according to

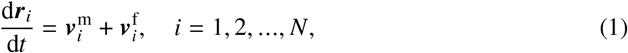

where 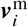 represents tangential motion arising from mechanical relaxation, and 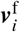 represents normal motion due to tissue formation.

The velocity 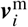 is constructed under the assumption of overdamped motion, appropriate for slow cellular dynamics in viscous environments (Purcell, 1977). In this regime, inertial effects are neglected and velocity is proportional to the net force acting on the boundary ***r***_*i*_ (*t*) such that

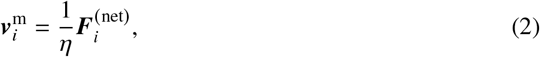

where *η* is the viscous drag coefficient. The net force is composed of three contributions (Figure 2b): two restoring forces, 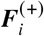 and 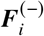, from the cells on either side, and a reaction force 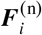 from the existing substrate. The restoring forces act along the cell bodies, with magnitudes determined by a prescribed force law *f* (*ℓ*_*i*_ (*t*)) which may be Hookean or nonlinear, and *ℓ*_*i*_ (*t*) = ∥ ***r***_*i*_ (*t*) − ***r***_*i*−1_(*t*) ∥ is the cell length (Murray et al., 2012; Murphy et al., 2019). he reaction force is defined to oppose the normal component of the restoring forces, thereby restricting mechanical relaxation to the tangent direction of the interface 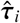. Consequently, the net force is given by the tangential projection

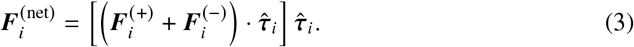

The velocity contribution 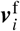, models the perpendicular displacement of cells in the direction 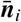 as they deposit ECM at the interface (Figure 2a). Compatibility considerations requires that the magnitude of this velocity contribution to satisfy

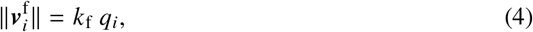

where *q*_*i*_ = 1/*ℓ*_*i*_ is the density of the *i*th cell (Buenzli, 2015; Alias and Buenzli, 2017; Hegarty-Cremer et al., 2021; Kuba et al., 2026). In this discrete framework, perpendicular displacements of the cell segments can cause them to overlap or separate, which leads to a disconnected interface. To combat this issue we implement a predictor-corrector algorithm to calculate 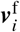 depending on whether neighboring segments intersect or separate after being displaced. A full description of this algorithm is described in detail in Kuba et al. (2026).

In addition to the mechanical and formation processes, the model incorporates four stochastic cell events:

- **Proliferation (Symmetric division)**: a cell divides into two active tissue-forming daughter cells that both lie on the tissue interface (Figure 2c).
- **Proliferation (Asymmetric division)**: a cell divides into two daughter cells where one remains an active tissue-forming cell and the other differentiates and embeds into the ECM (Figure 2d).
- **Apoptosis**: a cell dies and is then removed from the tissue interface (Figure 2e).
- **Differentiation**: a cell differentiates and embeds into the ECM (Figure 2f).

A key novelty of the framework is that it explicitly records the positions of differentiated cells (marked in red in Figure 2a,d,f) during asymmetric division and differentiation, enabling single-cell resolution comparisons between model predictions and experimental observations. Although asymmetric division and direct differentiation both generate differentiated cells, they represent distinct cellular mechanisms. While these processes are not easily distinguishable in static experimental observations, they can be independently controlled and analysed within the computational framework.

These events are assumed to occur over a sufficiently small time interval Δ*t*, such that at most one event takes place within [*t, t* + Δ*t*) (Baker et al., 2019; Murphy et al., 2020). Each cell may undergo proliferation (either symmetric division or asymmetric division), apoptosis, or differentiation with constant rates (probabilities per unit time) *P*_sym_, *P*_asym_, *A*, and *D*, respectively. Stochastic implementation proceeds by generating three random numbers from a uniform distribution to determine: (i) whether an event occurs, (ii) which event occurs, and (iii) which cell undergoes the event (Gelman et al., 2013; Baker et al., 2019; Murphy et al., 2020). Implementation details are provided in A.1 and open-source code is available at Kuba et al. (2026).

### 2.2 Connecting model parameters to experimental observations

To enable comparison between model-generated and experimental tissue compositions, model parameters were calibrated to reproduce experimentally measured differentiated cell densities. In the model, differentiated cells arise at the moving tissue interface through either direct differentiation (*D*) or asymmetric division (*P*_asym_), such that the total differentiation rate is

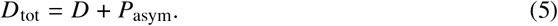

Following Buenzli (2015), experimentally consistent differentiated cell densities *ρ* are obtained by approximating

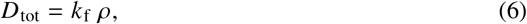

where *k*_f_ is the tissue formation rate. For each case study, experimentally estimated values of *ρ* and *k*_f_ (Section 2.3) were substituted into Eq. (6) to determine *D*_tot_.

To characterize the relative contribution of differentiation pathways, we define

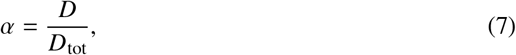

representing the fraction of differentiation occurring through direct differentiation. Since direct differentiation removes an active tissue-forming cell from the interface, increasing *α* reduces the density of active cells and influences interface morphology via the normal velocity formulation (Eq. (4)).

The evolution of the active tissue-forming cell population *N* is described by

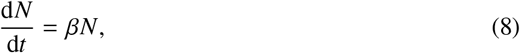

where *β* = *P*_sym_ − *A* − *D* is the net growth rate. Here, *β* > 0 corresponds to a proliferative population and *β* < 0 to a declining population. Together, *α* and *β* provide independent control over differentiation mode and population dynamics in the simulations

### 2.3 Experimental data

Previously published experimental measurements of differentiated cell density and extracellular matrix (ECM) formation rates were used to calibrate model parameters and ensure consistency between simulated and observed tissue compositions. Two experimental contexts were considered: (i) tissue engineering cultures within 3D-printed constructs, and (ii) bone formation within cortical pores.

For tissue engineering applications, data were drawn from in vitro studies of MC3T3-E1 osteoblastic cells cultured in thin 3D-printed porous scaffolds (Buenzli et al., 2020; Lanaro et al., 2021; Browning et al., 2021). In those experiments, cells were seeded within square pores of side length 200–600 μm and cultured for up to 28 days. Tissue growth was monitored using DAPI staining of cell nuclei and actin filaments, with ethidium homodimer staining indicating the absence of cell death (Buenzli et al., 2020). Images were acquired at multiple time points (4, 7, 10, 14, 18, and 28 days), from which differentiated cell density, cell orientation, tissue area coverage, and pore infill duration were quantified (Lanaro et al., 2021; Browning et al., 2021). Although orientation, coverage, and infill time varied with pore size, parameter estimation across all pores indicated that at late time (*t* > 14 days) the density of differentiated cells remain around *ρ* ≈ 0.003 cells/μm^2^ (Browning et al., 2021). The corresponding ECM formation rate was estimated as *k*_f_ ≈ 87.84 μm^2^/day (Kuba et al., 2026).

For cortical bone formation, matrix apposition rate (MAR) and osteocyte density (*ρ*) were obtained from previously published double-labeling experiments and imaging studies in animal models and human bone (Metz et al., 2003; Hannah et al., 2010; Power et al., 2012). Reported MAR values in human cortical bone range between 0.5–3 μm/day (Jones, 1974; Martin et al., 1998; Buenzli et al., 2014; Alias and Buenzli, 2018). To ensure consistency with this range (target ≈ 1.5 μm/day), the model parameter *k*_f_ was set to 30 μm^2^/day, assuming a typical osteoblast size of ∼ 20 μm and using Eq. (4). Consistent with reported estimates (Alias and Buenzli, 2018; Parfitt, 1983; Hannah et al., 2010; Power et al., 2012), osteocyte density was taken as *ρ* = 0.000625 cells/μm^2^ throughout the pore.

## 3 Results

We apply the computational model to two case studies: (i) tissue growth within 3D-printed scaffolds (Section 3.1) and (ii) bone formation within cortical pores (Section 3.2). In the tissue engineering study, we investigate how differentiation mechanisms, cell population dynamics, and mechanical properties of cells shape tissue growth patterns, interface morphology, and differentiated cell orientations. In the bone formation case study we examine how stochasticity influences the distribution of differentiated cells and the structural symmetry of the pore.

### 3.1 Tissue Engineering

To reflect the experimental setup described in Section 2.3, the tissue interface was initialized in a 300 μm × 300 μm square with *N* = 60 uniformly distributed cells of length 20 μm, approximating osteoblastic size. From Eq. (6), we estimated *D*_tot_ ≈ 0.26, and due to the absence of cell death we set *A* = 0. Since the fraction of differentiated cells (*α*) that emerge from direct differentiation and the net growth rate of the tissue-forming population (*β*) are difficult to measure experimentally, we investigated their effects by varying these parameters.

Figure 3a shows representative model-generated tissue compositions, with differentiated cells shown in red and the tissue interface indicated at the initial time (dark blue) and at 90% pore fill (blue). Within the newly formed tissue, differentiated cells vary in orientation and size but remain approximately uniformly distributed, despite the increasing density of tissue-forming cells as the interface evolves. To investigate the influence of differentiation modes, in Figure 3a we varied *α* and set *β* = 0 so that *P*_sym_, *P*_asym_, and *D* balance to maintain a consistent population of tissue-forming cells. Comparison with experimental observations (Figure 3b) indicates that *α* = 0.2 provides the closest agreement between model predictions and experiments, whereas larger *α* values yield a rougher and less consistent interface morphology. Figure 3c shows the effect of a dynamic tissue-forming cell population on the time required to achieve pore infilling. In these simulations, *α* was fixed at 0.2, and growing (*β* > 0), constant (*β* = 0) and shrinking (*β* < 0) populations were considered. As in Figure 3a, simulations were performed until 90% pore fill, and the normalized tissue area, averaged from 50 realizations, was plotted over time for each *β* value alongside the averaged experimental data for scaffolds of this size.

**Figure 3.**
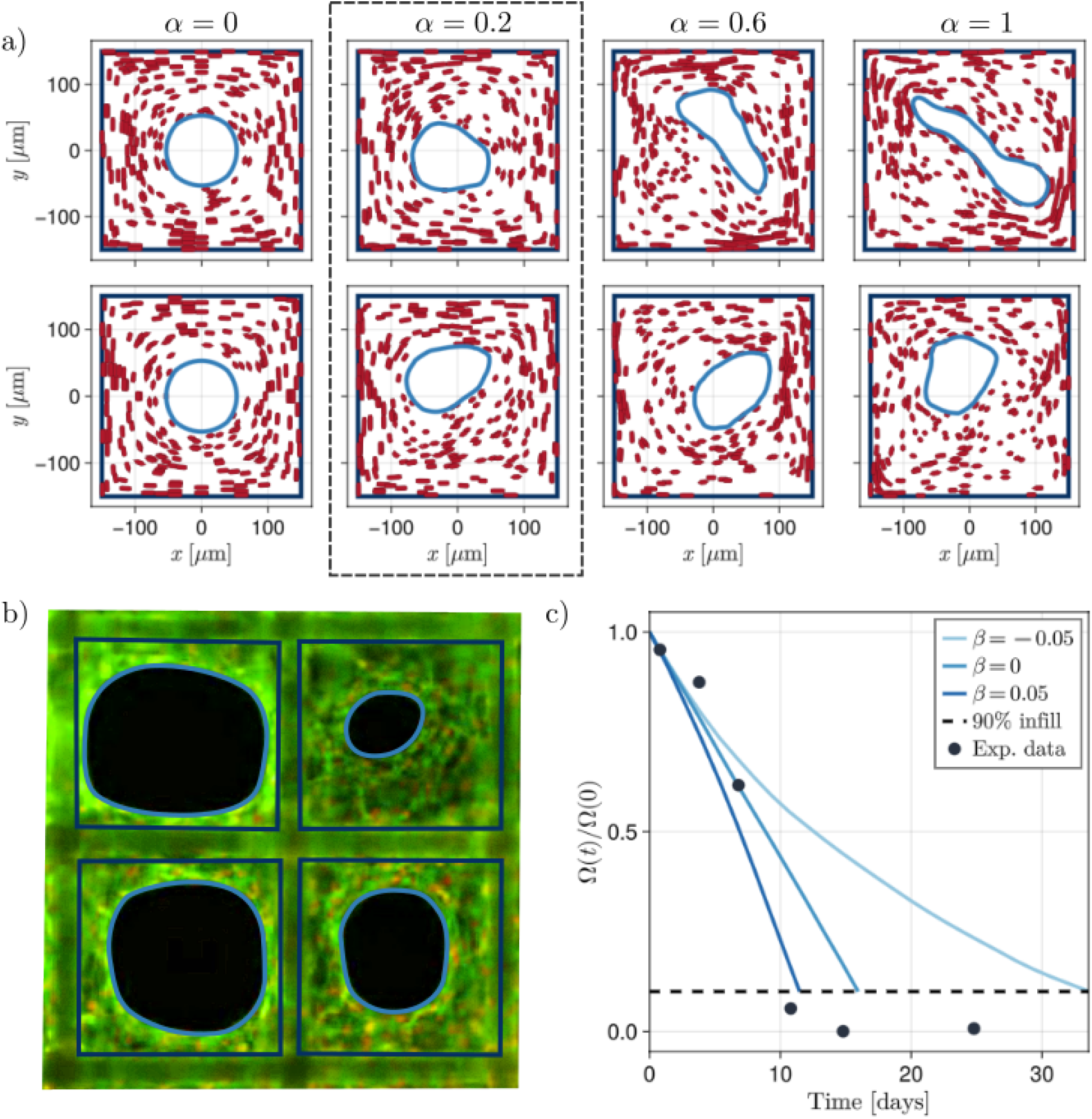
Example realisations of the tissue growth model for a series of control parameters *α* and *β* values. Simulations are initialised with a square geometry to emulate the cell culture experiments with *D*^tot^ = 0.003. (a) Realisations of the model for different values of *α* with *β* = 0. (b) Example images of experiments cultured withing 300 μm × 300 μm, reproduced with permissions from Buenzli et al. (2020). (c) Solid lines are the average normalized area of void space over time for different values of *β* with *α* = 0.2, and black points are averages estimated from experiments cultured in scaffolds of the same size. The simulated averages were generated from from a sample size of 50 realizations for each value of *β*.

Figure 4 examines the influence of cell mechanical stiffness on differentiated cell orientation. Stiffness levels were categorized as low (*k* = 5 N/μm), intermediate (*k* = 100 N/μm), and high (*k* = 500 N/μm). Representative realizations and corresponding rose plots show that orientation distributions are initially anisotropic across all stiffness levels (*t* = 4 days), but anisotropy decreases over time for lower stiffness values. Comparison of cumulative orientation distributions with experimental measurements (Lanaro et al., 2021) indicates improved agreement at lower stiffness, although the associated interface morphologies do not reproduce the experimentally observed rounding (Figure 5).

**Figure 4.**
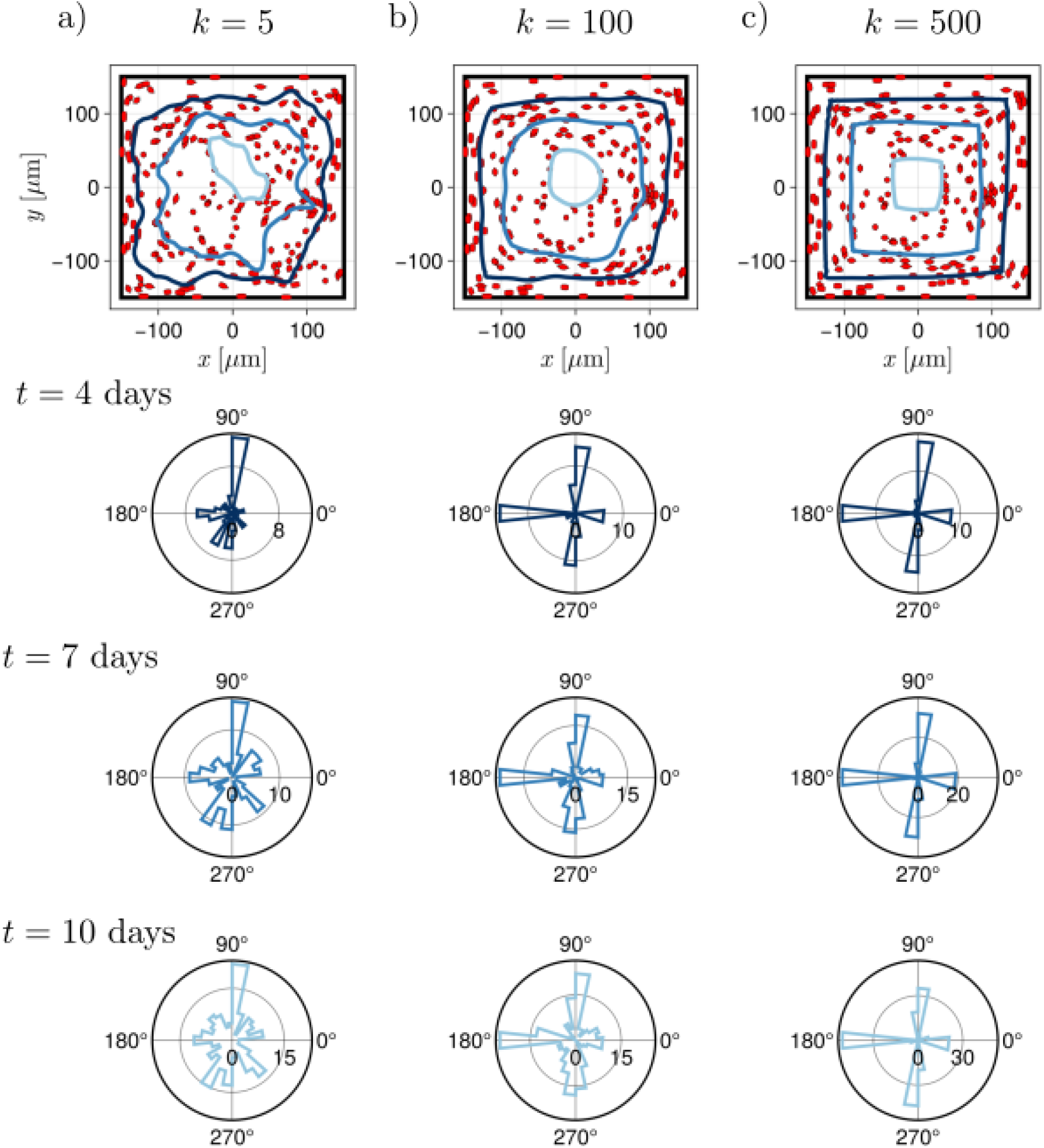
Simulations of the stochastic model showing the effect of cell stiffness on embedded cell orientations. Each column shows a single realization for (a) low (*k* = 5 N / μm) (b) intermediate (*k* = 100 N / μm) and (c) high (*k* = 500 N / μm) stiffness. The top row displays tissue growth with differentiated cells (red) and the interface at *t* = 0, 5, and 10 days, using a color gradient from dark blue (early) to light blue (late). Subsequent rows show the frequency of cell orientations measured relative to the x-axis and binned into 24 intervals over [0^◦^, 360^◦^).

**Figure 5.**
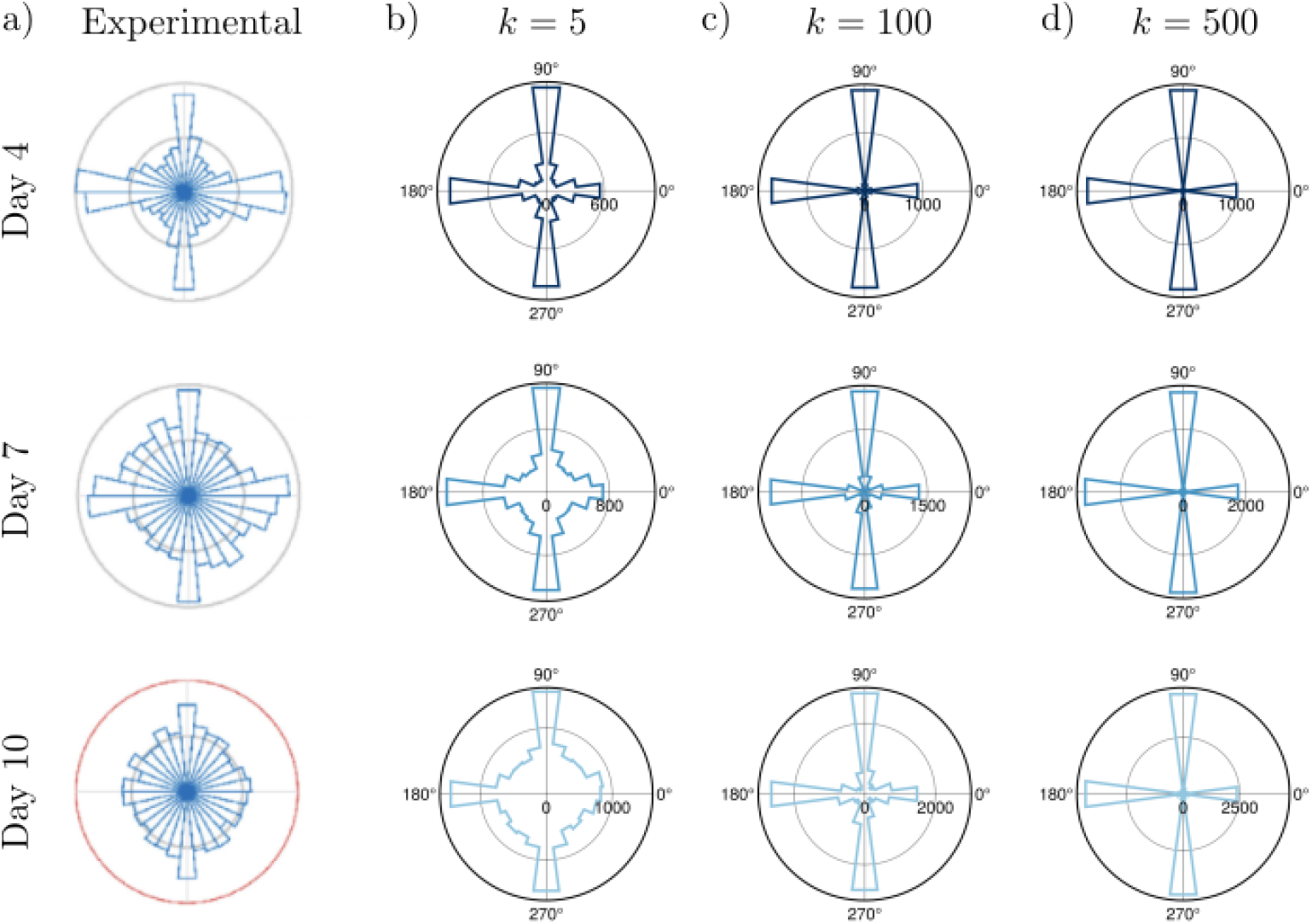
Comparison of experimental and simulated cell orientation distributions. (a) Measured cell orientations from cell culture experiments on days 4, 7, and 10 in a 300 μm × 300 μm 3D-printed square porous scaffold reported by Lanaro et al. (2021) (reproduced with permissions). (b-d) Cumulative cell orientation frequencies from 100 realizations for low (*k* = 5 N / μm), intermediate (*k* = 100 N / μm) and high (*k* = 500 N / μm) cell stiffness at simulation times that correspond to the experimental data.

### 3.2 Bone formation

To model bone tissue formation within irregular resorption cavities, the tissue interface was initialized as a deterministically perturbed circle. Specifically, *N* values were sampled from a uniform distribution and smoothed with respect to the angular coordinate *θ* ∈ [0, 2*π*) using the *loess* function to generate spatially correlated perturbations (Cleveland and Grosse, 1991). The resulting smoothed values *u*_*i*_ ∈ [0, 1] were scaled by a perturbation amplitude 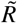 and added to a circle of radius *R*_0_ such that, at *t* = 0,

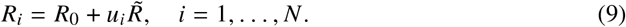

Experimental evidence indicates that bone formation initiates once osteoblast density reaches a threshold, at which point proliferation ceases (Lassen et al., 2017). Accordingly, proliferation rates were set to zero (*P*_sym_ = *P*_asym_ = 0), so differentiation occurred solely through direct differentiation, yielding *D* = *k*_f_*ρ* ≈ 0.019. The apoptosis rate *A* was chosen to be lower than *D* to prevent rapid cell loss.

Figure 6b shows a representative realization for an artificial resorption cavity, with osteocytes marked in black and growth regions shaded according to formation time, consistent with experimental images (Figure 6a). Osteocyte density was quantified within regions formed over a time interval of *T* = 7 days, corresponding to lamellar deposition in cortical bone. Summary statistics (Figure 6c) demonstrate that local osteocyte density remains consistent and closely matches the prescribed density *ρ*, despite variation in region shape and size over time.

**Figure 6.**
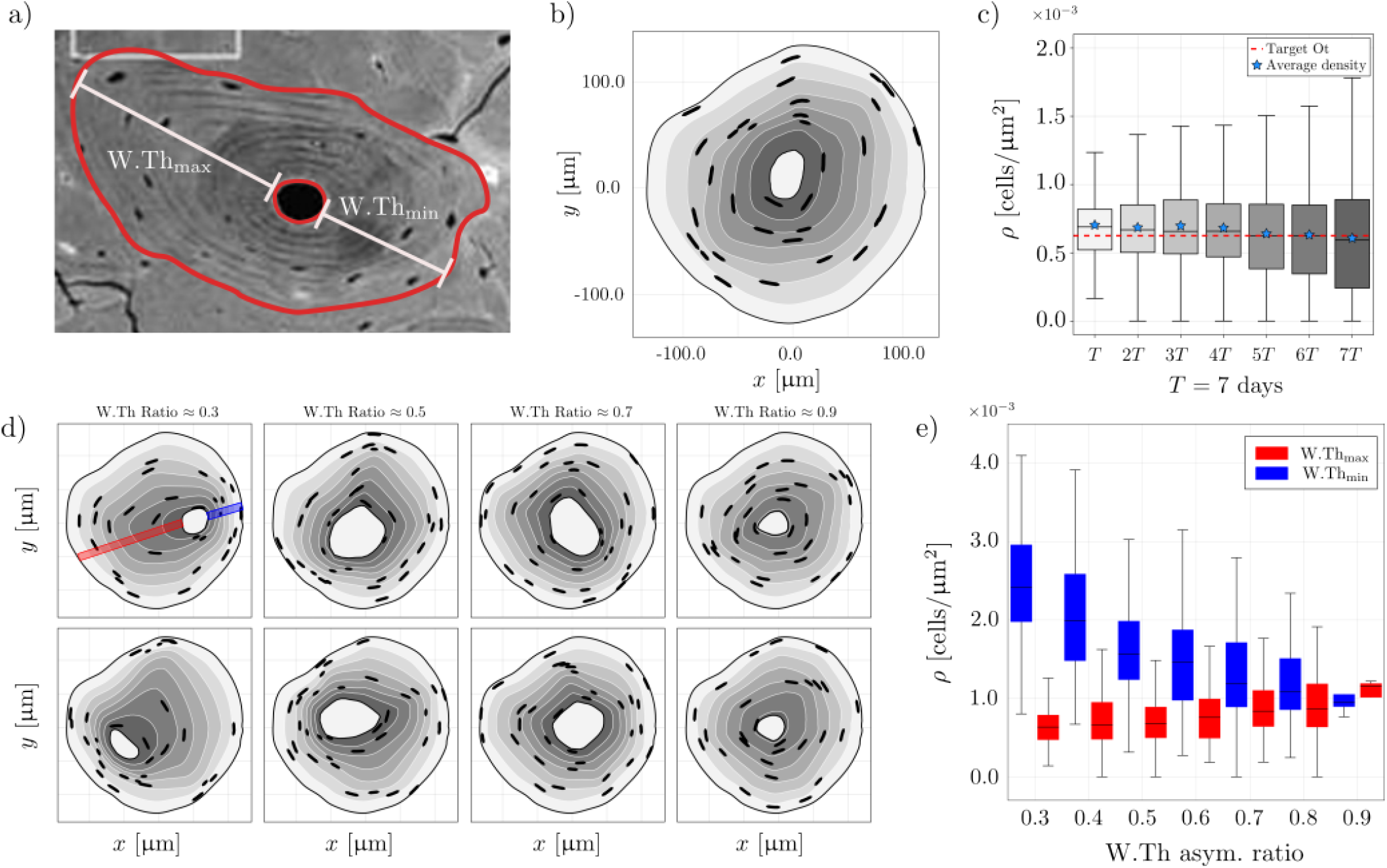
Model simulations of bone formation in an artificial osteon. (a) Experimental image illustrating the measurement of W.Th_max_ and W.Th_min_ used to compute the W.Th_asym_ ratio. (b) Representative model realization in an artificial osteon, with osteocytes marked in black. Pseudo-lamellae formed over a time interval *T* = 7 days are shaded according to formation time. (c) Box plots of embedded cell densities within wall regions formed over *T* days from 500 realizations. Shading corresponds to the wall regions illustrated in (a). (d) Example model realizations spanning a range of W.Th_asym_ values. (e) Summary statistics of embedded cell densities measured in rectangular regions near W.Th_max_ (blue) and W.Th_min_ (red), grouped by asymmetry ratio bins. Example sampling regions are highlighted in the top-left panel of (d). In all model realizations death rate of cells is *A* = 0.01 and total differentiation rate is *D*_tot_ = *k*_f_ *ρ* where *ρ* = 0.000625 cells/μm^2^.

Although the realization in Figure 6b shows symmetric pore closure, cortical pores may also develop asymmetric structures. Asymmetry was quantified using the wall thickness asymmetry ratio W.Th_asym_, defined using the pair of opposing wall thicknesses with the greatest disparity (Figure 6a), where wall thickness is measured as the distance between the inner and outer pore boundaries (Hegarty-Cremer et al., 2024):

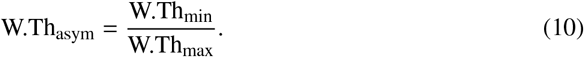

Figure 6d shows representative realizations classified by W.Th_asym_, and Figure 6e summarizes osteocyte densities in the wall regions used to compute this ratio. These results indicate that osteocyte density is higher near thinner walls, and the most frequently observed asymmetries fall within the range W.Th_asym_ ∈ [0.6, 0.7).

## 4 Discussion

Experimental evidence indicates that biological tissue growth is strongly regulated by substrate geometry and mechanobiological cues (Nelson et al., 2005; Rumpler et al., 2008; Bidan et al., 2012; Ladoux and Mége, 2017; Callens et al., 2020). In confined geometries, spatial constraints cause tissue-forming cells to crowd or spread during growth, generating mechanical stresses that in turn influence cellular processes such as proliferation and differentiation (Nelson et al., 2005; Werner et al., 2017; Gudipaty et al., 2017). However, disentangling the mechanistic interplay between geometry, mechanics, and cellular processes in tissue growth remains challenging when relying solely on experimental tissue samples, even though such samples contain fingerprints of the processes that shaped them. To address this limitation, we propose a computational model of tissue growth that generates synthetic tissues with single-cell resolution, enabling direct control of governing mechanisms and systematic comparison with experimental data to isolate their individual contributions.

During tissue growth, cells may undergo proliferation, differentiation, and apoptosis, with the relative contributions of these processes varying across tissue types. Histological staining can reveal the presence or absence of certain processes, but it cannot resolve the specific processes that occurred during growth. In tissue engineering experiments (Figure 1a), DAPI staining indicates that there is a substantial amount of cell proliferation during growth, while ethidium homodimer staining indicates the absence of cell death. Model simulations indicate that, given the incorporated mechanisms, closer agreement with experimental observations and smoother, more consistent tissue interfaces are achieved when the majority of differentiated cells arise from asymmetric division (*α* = 0.2) rather than direct differentiation (Figure 3a). In addition, comparison of normalized tissue area over time indicates that an increasing population of tissue-forming cells more accurately captures late-stage growth dynamics (Figure 3c), consistent with previous quantitative and modeling studies (Buenzli et al., 2020; Browning et al., 2021).

In these tissue engineered experiments, differentiated cells are assumed to become quiescent once embedded within the ECM. Consequently, differentiated cell orientations provide a record of the position and shape of the tissue interface at the time of differentiation. Previous modeling studies of evolving tissue interfaces have demonstrated a relationship between interface morphology and the strength of mechanical interactions between cells (Alias and Buenzli, 2017; Kuba et al., 2026). Consistent with this, model simulations show that intermediate cell stiffness produces tissue interfaces that more closely resemble the experimentally observed rounding (Figure 4), whereas low stiffness yields orientation distributions with reduced anisotropy that better match experimental measurements (Lanaro et al., 2021) (Figure 5). This discrepancy suggests that additional mechanisms may contribute to tissue growth mechanics beyond those captured by cell–cell mechanical interactions alone.

A plausible contributor to these additional mechanisms is surface tension. In their experimental study, Lanaro et al. (2021) observed groups of cells spanning perpendicular fibers and proposed that cells actively seek nearby attachment points to maintain surface tension during growth. This behavior is consistent with purse–string–based closure mechanisms observed in wound healing (Ravasio et al., 2015) and with evidence that surface tension plays a central role in tissue organization and mechanical regulation (Bidan et al., 2016). In the current model, surface tension–related forces acting perpendicular to the interface are not represented due to the definition of the reaction force 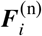. These findings underscore the importance of incorporating surface tension–mediated effects into tissue growth models, as neglecting them may obscure key mechanical contributions to interface evolution. Although this case study focused on square geometries, the same analysis can be extended to other pore shapes (Figure 7) to further examine the role of surface tension in tissue growth.

**Figure 7.**
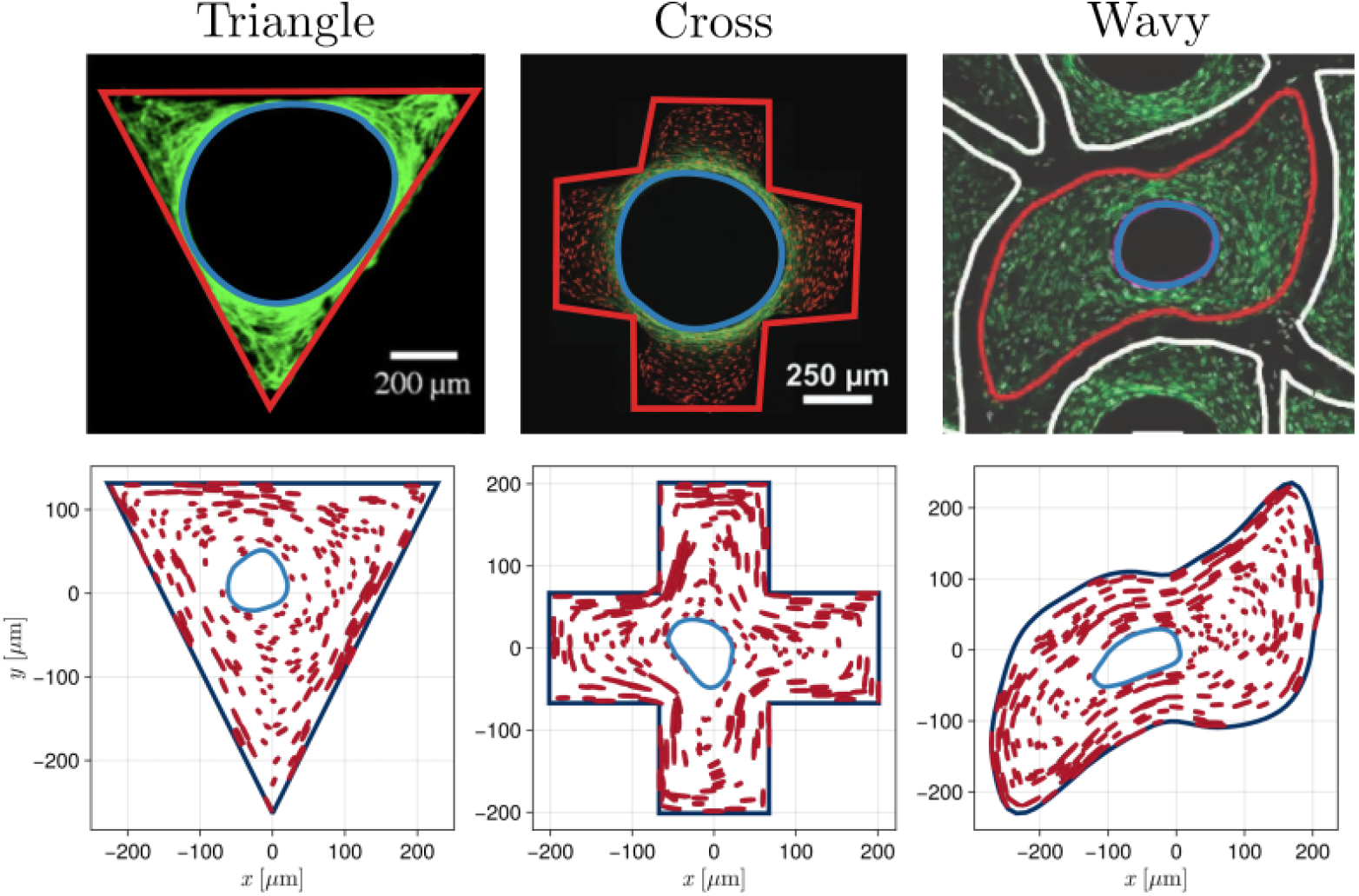
Example realisations of the tissue growth model in different initial geometries. The geometries are motivated by previously conducted cell culture experiments of Rumpler et al. (2008) (triangle), Bidan et al. (2013) (cross), and VandenHeuvel et al. (2023) (wavy).

In Section 3.2, we investigated how stochasticity influences the spatial distribution of differentiated cells and the resulting growth patterns during bone formation within cortical pores. In this setting, a finite population of osteoblasts deposits new matrix at the pore surface without proliferation, differentiating into osteocytes that become embedded within the mineralized tissue. Experimental studies report that osteocyte density decreases with distance from the initial pore boundary (Power et al., 2012), although differentiation rates cannot be directly measured. Consistent with previous modeling work (Buenzli, 2015), our simulations show that osteocyte density within newly formed lamellar regions remains consistent with the prescribed constant value *ρ*, despite variations in regional shape and size (Figure 6b,c). This suggests that dynamic differentiation rates could be inferred from single experimental snapshots by quantifying osteocyte densities within lamellae and rearranging Eq. (6), providing a basis for incorporating curvature-, stress-, or property-dependent differentiation rules (e.g., *D*(*κ, σ*, …)) into the modeling framework.

Cortical bone formation occurs around Haversian canals whose positions are often asymmetric (Robling, 1998; Schnitzler and Mesquita, 2013; Cooke et al., 2022), motivating investigation into the mechanisms underlying such asymmetries. Previous modeling studies have attributed asymmetric growth to biological heterogeneities, such as spatial variations in matrix deposition rates (Hegarty-Cremer et al., 2024). In contrast, our analysis (Figure 6d,e) demonstrates that asymmetries can arise purely from stochastic differentiation and apoptosis of bone-forming cells. The discrete formulation of the model captures density fluctuations associated with these events, leading to locally reduced normal velocities (Eq. (4)) and emergent interface irregularities. These findings suggest that at least part of the variability observed in cortical pore morphology may result from intrinsic stochastic cellular processes, although further experimental validation is required.

## 5 Conclusions and future work

Experimental evidence demonstrates that tissue growth is governed by the interplay between geometry, mechanics, and biological processes (Nelson et al., 2005; Rumpler et al., 2008; Bidan et al., 2012; Ladoux and Mége, 2017; Callens et al., 2020). Disentangling these mechanisms experimentally remains challenging, as mechanical properties are difficult to measure (Tee et al., 2009) and tissue samples provide only static snapshots of a dynamic process. However, the material deposited behind the advancing tissue interface retains structural signatures of the processes that shaped tissue architecture. Here, we introduced a computational model that simulates tissue growth at single-cell resolution and generates synthetic tissue compositions directly comparable with experimental observations. Through two case studies, we demonstrated that the framework can (i) identify the contribution of specific mechanisms to tissue morphology, (ii) estimate dynamic rates from single-time-point data, and (iii) generate testable hypotheses for the origins of stochastic growth patterns. These results highlight the utility of model-generated tissue compositions as a complementary tool for investigating tissue growth dynamics and enabling in silico hypothesis testing.

Future extensions could focus on tailoring the framework to specific tissue types by incorporating biologically informed process rules. For example, in bone formation, differentiation rates could be extended to depend on interface curvature or mechanical stress and calibrated using experimental estimates. Similar extensions could be applied to proliferation, apoptosis, and the tissue formation rate *k*_f_, consistent with previous studies in which *k*_f_ depended on interface radius or porosity (Alias and Buenzli, 2018), and cell event rates depended on cell length (Murphy et al., 2020). Parameter estimation could then be performed by comparing synthetic and experimental datasets using statistical inference techniques. Additionally, the model’s ability to track embedded cell positions enables spatial statistical analyses, such as point process or pair-correlation methods, providing a quantitative framework for testing competing mechanistic hypotheses and guiding future experimental design.

## Acknowledgements

This researchwas supported by the Australian Research Council (DP190102545, DP230100025). SK and PRB acknowledge support from the Max Planck Queensland Centre for the Materials Science of Extracellular Matrices. We thank Brenna Devlin and Maria A. Woodruff for providing the experimental image presented in Figure 1a.

## Appendix A Mathematical Model

### A.1 Implementation of stochastic cellular processes in mathematical model

To simplify notation, we write the probability that the *i*th cell proliferates is *P*_sym *i*_ (symmetric division). Similarly for asymmetric proliferation, apoptosis, and differentiation, we write the probabilities as *P*_asym *i*_, *A*_*i*_, and *D*_*i*_. The algorithm at each numerical time step Δ*t* has been implemented as follows:

1. Generate *p*_1_, *p*_2_, *p*_3_ from a uniform distribution in [0, 1].
2. If 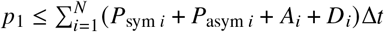, where *N* is the total number of cells that form the interface, then one of the cells is experiencing one of the cellular processes over the time interval [*t* + Δ*t*). Otherwise continue to the next simulation time step.
3. Using *p*_2_, we determine which cellular process occurred under the following conditions:
  - If 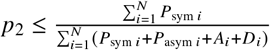, then the occurring cellular process is proliferation.
  - Else if 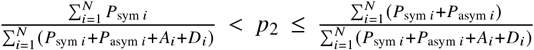 then the occurring cellular process is asymmetric proliferation.
  - Else if 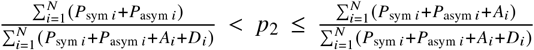 then the occurring cellular process is apoptosis.
  - Else the occurring cellular process is differentiation.
4. Using *p*_3_, we determine which cell experienced the cellular process. For example, assuming that the previous step resulted in proliferation (symmetric division), we assert that the *j* th cell proliferates when the conditions below are met,

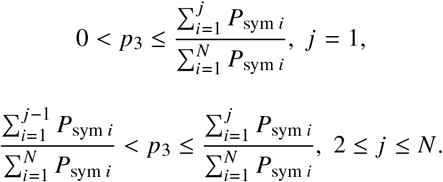 A similar process is followed in the case that the previous step results in apoptosis, embedment, and simultaneous proliferation and embedment.
5. Based on cell event, we either add or remove cell boundaries from the interface, linearly interpolating the boundary nodes over the region where the *j* th cell spanned as described in Section 2.1.

## References

Poujade, M., Grasland-Mongrain, E., Hertzog, A., Silberzan, P., Collective migration of an epithelial monolayer in response to a model wound., Cell Biology, 2007; 104: 15988–15993. doi: 10.1073/pnas.0705062104.

Rolli, C.G., Nakayama, H., Yamaguchi, K., Spatz, J.P., Kemkemer, R., Nakanishi, J., Switchable adhesive substrates: Revealing geometry dependence in collective cell behavior., Biomaterials, 2012; 33: 2409–2418. doi: 10.1016/j.biomaterials.2011.12.012.

Martin, R.B., Burr, D.B., Sharkey, N.A., Fyhrie, D.P. Skeletal Tissue Mechanics., New York: springer, 1998.

Lowengrub, J.S., Frieboes, H.B., Jin, F., Chuang, Y-L., Li, X., Macklin, P., Wise, S.M., Cristini, V., Nonlinear modelling of cancer: bridging the gap between cells and tumours., Nonlinearity, 2009; 23: R1. doi: 10.1088/0951-7715/23/1/R01.

Wozniak, P., El haj, A.J., Tissue Engineering Using Ceramics and Polymers, Chapter 14 - Bone regeneration and repair using tissue engineering., Woodhead Publishing, 2007.

Hollister, S.J., Porous scaffold design for tissue engineering, Nature Materials, 2005; 4: 518– 524. doi: 10.1038/nmat1421.

O’Brien, F.J., Biomaterials & scaffolds for tissue engineering, Materials Today, 2011; 14: 88–95. doi: 10.1016/S1369-7021(11)70058-X.

Bidan, C.M., Kommareddy, K.P., Rumpler, M., Kollmannsberger, P., Fratzl, P., Dunlop, J.W.C., Geometry as a Factor for Tissue Growth., Advanced Healthcare Materials, 2013; 2: 186–194. doi: 10.1002/adhm.201200159.

Rumpler, M., Woesz, A., Dunlop, J.W.C, van Dongen, J.T, Fratzl, P., The effect of geometry on three-dimensional tissue growth., Journal of The Royal Society Interface, 2008; 27: 1173– 1180. doi: 10.1098/rsif.2008.0064.

Bidan, C., Kommareddy, K., Rumpler, M., Kollmannsberger, P., Bréchet, Y., Fratzl, P., Dunlop, J., How Linear Tension Converts to Curvature: Geometric Control of Bone Tissue Growth., PloS one, 2012; 7: e36336. doi: 10.1371/journal.pone.0036336.

Schamberger, B., Ziege, R., Anselme, K., Ben Amar, M., Bykowski, M., Castro, A.P.G., Cipitria, A., Coles, R.A., Dimova, R., Eder, M., Ehrig, S., Escudero, L.M., Evans, M.E., Fernandes, P.R., Fratzl, P., Geris, L., Gierlinger, N., Hannezo, E., Iglič, A., Kirkensgaard, J.J.K., Kollmannsberger, P., Kowalewska, Ł., Kurniawan, N.A., Papantoniou, I., Pieuchot, L., Pires, E.F., Roschger, A., Bidan, C.M., Dunlop, J.W.C., Curvature in Biological Systems: Its Quantification, Emergence, and Implications across the Scales., Advanced Materials, 2023; 35: 2206110. doi: 10.1002/adma.202206110.

Callens, S.J.P., Uyttendaele, R.J.C., Fratila-Apachitei, L.E., Zadpoor, A.A., Substrate curvature as a cue to guide spatiotemporal cell and tissue organization., Biomaterials, 2020; 232: 119739. doi: 10.1016/j.biomaterials.2019.119739.

McNamara, L.E., McMurray, R.J., Dalby, M.J., Nanotopographical Control of Stem Cell Differentiation., Journal of Tissue Engineering, 2010; 1: 120623. doi: 10.4061/2010/120623.

Dobbenga, S., Fratila-Apachitei, L.E., Zadpoor, A.A., Nanopattern-induced osteogenic differentiation of stem cells – A systematic review., Acta Biomaterialia, 2016; 46: 3–14. doi: 10.1016/j.actbio.2016.09.031.

Nelson, C.M., Jean, R.P., Tan, J.L., Liu, W.F., Sniadecki, N.J., Spector, A.A., Chen, C.S., Emergent patterns of growth controlled by multicellular form and mechanics., Proceedings of the National Academy of Sciences, 2005; 102: 11594–11599. doi: 10.1073/pnas.0502575102.

Discher, D.E., Janmey, P., Wang, Y-L., Tissue Cells Feel and Respond to the Stiffness of Their Substrate., American Association for the Advancement of Science, 2005; 310: 1139–1143. doi: 10.1126/science.1116995.

Ladoux, B., Mége, R.M., Mechanobiology of collective cell behaviours., Nature Reviews Molecular Cell Biology, 2017; 18: 743–757. doi: 10.1038/nrm.2017.98.

Lim, C.T., Bershadsky, A., Sheetz M.P., Mechanobiology., Journal of The Royal Society Interface, 2010; 7: S291–S293. doi: 10.1098/rsif.2010.0150.focus.

Alberts, B., Johnson, A., Lewis, J., Raff, M., Roberts, K., Walter, P., Molecular Biology of the Cell, 4th edition., New York: Garland Science, 2002.

Kurniawan, N.A., Bouten, C.V.C., Mechanobiology of the cell–matrix interplay: Catching a glimpse of complexity via minimalistic models, Extreme Mechanics Letters, 2018; 20: 59–64. doi: 10.1016/j.eml.2018.01.004.

Skedros, J.G., Holmes, J.L., Vajda, E.G., Bloebaum, R.D., Cement lines of secondary osteons in human bone are not mineral-deficient: New data in a historical perspective., The Anatomical Record, 2005; 286: 781–803. doi: 10.1002/ar.a.20214.

Roschger, P., Paschalis, E.P., Fratzl, P., Klaushofer, K., Bone mineralization density distribution in health and disease., Bone, 2008; 42: 456–466. doi: 10.1016/j.bone.2007.10.021.

Vatsa, A., Breuls, R.G., Semiens, C.M., Salmon, P.L., Smit, T.H., Klein-Nulend, J., Osteocyte morphology in fibula and calvaria — Is there a role for mechanosensing?, Bone, 2008; 43: 452–458. doi: 10.1016/j.bone.2008.01.030.

Ascenzi, M., Gills, J., Lomovstev, A., Orientation of collagen at the osteocyte lacunae in human secondary osteons., Journal of Biomechanics, 2008; 41: 3426–3435. doi: 10.1016/j.jbiomech.2008.09.010.

van Hove, R.P., Nolte, P.A., Vatsa, A., Semiens, C.M., Salmon, P.L., Smit, T.H., Klein-Nulend, J., Osteocyte morphology in human tibiae of different bone pathologies with different bone mineral density — Is there a role for mechanosensing?, Bone, 2009; 45: 321–329. doi: 10.1016/j.bone.2009.04.238.

Britz, H.M., Carter, Y., Jokihaara, J., Leppanen, O.V., Jarvinen, T.L.N., Belev, G., Cooper, D.M.L., Prolonged unloading in growing rats reduces cortical osteocyte lacunar density and volume in the distal tibia., Bone, 2012; 51: 913–919. doi: 10.1016/j.bone.2012.08.112.

Mader, K.S., Schneider, Philipp, S., Muller, R., Stampanoni, M., A quantitative framework for the 3D characterization of the osteocyte lacunar system., Bone, 2013; 57: 142–154. doi: 10.1016/j.bone.2013.06.026.

Lanaro, M., Mclaughlin, M.P., Simpson, M.J., Buenzli, P.R., Wong, C.S., Allenby, M.C., Woodruff, M.A., A quantitative analysis of cell bridging kinetics on a scaffold using computer vision algorithms., Acta Biomaterialia, 2021; 136: 429–440. doi: 10.1016/j.actbio.2021.09.042.

Ruffoni, D., Fratzl, P., Roschger, P., Klaushofer, K., Weinkamer, R., The bone mineralization density distribution as a fingerprint of the mineralization process., Bone, 2007; 40: 1308– 1319. doi: 10.1016/j.bone.2007.01.012.

Buenzli, P.R., Osteocytes as a record of bone formation dynamics., Journal of Theoretical Biology, 2015; 364: 418–427. doi: 10.1016/j.jtbi.2014.09.028.

Cox, B.N., Purohit, P.K., White, S.N. Strain fields and solitary strain waves as determining factors for the cross-sectional geometry of mouse incisor enamel., Journal of the Mechanics and Physics of Solids, 2015; 193: 105840. doi: 10.1016/j.jmps.2024.105840.

Guyot, Y., Papantoniou, I., Chai, Y.C., Van Bael, S., Schrooten, J., Geris, L., A computational model for cell/ECM growth on 3D surfaces using the level set method: a bone tissue engineering case study., Biomechanics and Modeling in Mechanobiology, 2014; 13: 1361–1371. doi: 10.1007/s10237-014-0577-5.

Callens, S.J.P., Fan, D., van Hengel, I.A.J., Minneboo, M., Díaz-Payno, P.J., Stevens, M.M., Fratila-Apachitei, L.E., Zadpoor, A.A., Emergent collective organization of bone cells in complex curvature fields., Nature Communications, 2023; 14: 855. doi: 10.1038/s41467-023-36436-w.

Ambrosi, D., Guana, F., Stress-Modulated Growth., Mathematics and Mechanics of Solids, 2007; 12: 319–342. doi: 10.1177/1081286505059739.

Dunlop, J.W.C., Fischer, F.D., Gamsjäger, E., Fratzl, P., A theoretical model for tissue growth in confined geometries., Journal of the Mechanics and Physics of Solids, 2010; 58: 1073–1087. doi: 10.1016/j.jmps.2010.04.008.

Goriely, A., The Mathematics and Mechanics of Biological Growth., Springer New York, NY, 2017.

Joly, P., Duna, G.N., Schone, M., Welzel, P.B., Fruedenberg, U., Werner, C., Petersen, A., Geometry-Driven Cell Organization Determines Tissue Growths in Scaffold Pores: Consequences for Fibronectin Organization., PLOS One, 2013; 8: e73545. doi: 10.1371/journal.pone.0073545.

Gahffarizadeh, A., Heiland, R., Friedman, S.H., Mumenthaler, S.M., Macklin, P., PhysiCell: An open source physics-based cell simulator for 3-D multicellular systems., PLOS Computational Biology, 2018; 14: e1005991. doi: 10.1371/journal.pcbi.1005991.

Alias, M.A., Buenzli, P.R., Modeling the Effect of Curvature on the Collective Behavior of Cells Growing New Tissue., Biophysical Journal, 2017; 112: 193–204. doi: 10.1016/j.bpj.2016.11.3203.

Hegarty-Cremer, S.G.D., Simpson, M.J., Andersen, T.L., Buenzli, P.R., Modelling cell guidance and curvature control in evolving biological tissues., Journal of Theoretical Biology, 2021; 520: 110658. doi: 10.1016/j.jtbi.2021.110658.

Kuba, S., Simpson, M.J., Buenzli, P.R., A mathematical model of curvature controlled tissue growth incorporating mechanical cell interactions, 2026; BioRxiv. doi: 10.64898/2026.03.10.710423.

Baker, R.E., Parker, A., Simpson, M.J., A free boundary model of epithelial dynamics., Journal of Theoretical Biology, 2019; 481: 61–74. doi: 10.1016/j.jtbi.2018.12.025.

Murphy, R.J., Buenzli, P.R., Baker, R.E., Simpson, M.J., A one-dimensional individual-based mechanical model of cell movement in heterogeneous tissues and its coarse-grained approximation., Proceedings of the Royal Society A, 2019; 475: 20180838. doi: 10.1098/rspa.2018.0838.

Buenzli, P.R., Kuba, S., Murphy, R.J., Simpson, M.J., Mechanical cell interactions on curved interfaces., Bulletin of Mathematical Biology, 2025; 87: 29. doi: 10.1007/s11538-024-01406-w.

Brown, P.J., Green, E.F., Binder, B.J., Osborne, J.M., Competing mechanisms for the buckling of an epithelial monolayer identified using multicellular simulation., Mathematical Biosciences, 2025; 380: 109367. doi: 10.1016/j.mbs.2024.109367.

Murray, P.J., Edwards, C.M., Tindall, M.J., Maini, P.K., From a discrete to a continuum model of cell dynamics in one dimension., Physical Review E, 2009; 80: 031912. doi: 10.1103/PhysRevE.80.031912.

Murray, P.J., Edwards, C.M., Tindall, M.J., Maini, P.K., Classifying general nonlinear force laws in cell-based models via the continuum limit., Physical Review E, 2012; 85: 021921. doi: 10.1103/PhysRevE.85.021921.

Gelman, A., Carlin, J.B., Stern, H.S., Dunson, D.B., Vehtari, A., Rubin, D.B., Bayesian Data Analysis, Third Edition, Chapman and Hall/CRC, 2013.

Cleveland, W.S., Grosse, E., Computational methods for local regression, Statistics and Computing, 1991; 1(1), 47–62. doi: 10.1007/BF01890836.

Purcell, E.J., Life at low Reynolds number, American Journal of Physics 1977; 45: 3–11. doi: 10.1142/9789814434973_0004.

Murphy, R.J., Buenzli, P.R., Baker, R.E., Simpson, S.J., Mechanical Cell Competition in Heterogeneous Epithelial Tissues., Bulletin of Mathematical Biology, 2020; 82: 130. doi: 10.1007/s11538-020-00807-x.

Buenzli, P.R., Lanaro, M., Wong, C.S., McLaughlin, M.P., Allenby, M.C., Woodruff, M.A., Simpson, M.J., Cell proliferation and migration explain pore bridging dynamics in 3D printed scaffolds of different pore size., Acta Biomaterialia, 2020; 114: 285–295. doi: 10.1016/j.actbio.2020.07.010.

Browning, A.P., Maclaren, O.J., Buenzli, P.R., Lanaro, M., Allenby, M.C., Woodruff, M.A., Simpson, M.J., Model-based data analysis of tissue growth in thin 3D printed scaffolds., Journal of Theoretical Biology, 2021; 528: 110852. doi: 10.1016/j.jtbi.2021.110852.

Parfitt, A.M., The physiological and clinical significance of bone histomorphometric data., In Recker R.R. (Ed.), Bone histomorphometry: Techniques and interpretation., 1983.

Alias, M.A., Buenzli, P.R., Osteoblasts infill irregular pores under curvature and porosity controls., Biomechanics and Modeling in Mechanobiology, 2018; 17: 1357–1371. doi: 10.1007/s10237-018-1031-x.

Metz, L.N., Martin, R.B., Turner, A.S., Histomorphometric analysis of the effects of osteocyte density on osteonal morphology and remodeling., Bone, 2003; 33: 753–759. doi: 10.1016/S8756-3282(03)00245-X.

Hannah, K.M., Thomas, C.D.L., Clement, J.G., Peele, A.G., Bimodal distribution of osteocyte lacunar size in the human femoral cortex as revealed by micro-CT., Bone, 2010; 47: 866–871. doi: 10.1016/j.bone.2010.07.025.

Power, J., Doube, M., van Bezooijen, R.L., Loveridge, N., Reeve, J., Osteocyte recruitment declines as the osteon fills in: Interacting effects of osteocytic sclerostin and previous hip fracture on the size of cortical canals in the femoral neck., Bone, 2012; 50: 1107–1114. doi: 10.1016/j.bone.2012.01.016.

Jones S.P., Secretory territories and rate of matrix production of osteoblasts., Calcified Tissue Research, 1974; 14: 309–315. doi: 10.1007/BF02060305.

Martin, R.B., Burr, D.B., Sharkey, N.A., Skeletal Tissue Mechanics., Springer New York, NY, 1998.

Buenzli, P.R., Pivonka, P., Smith, D.W., Bone refilling in cortical basic multicellular units: insights into tetracycline double labelling from a computational model., Biomechanics and Modeling in Mechanobiology, 2014; 13: 185–204. doi: 10.1007/s10237-013-0495-y.

Bidan, C.M., Kollmannsberger, P., Gering, V., Ehrig, S., Joly, P., Petersen, A., Vogel, V., Fratzl, P., Dunlop, J.W.C., Gradual conversion of cellular stress patterns into pre-stressed matrix architecture during in vitro tissue growth., Journal of The Royal Society Interface, 2016; 13: 20160136. doi: 10.1098/rsif.2016.0136.

VandenHeuvel, D.J., Devlin, B.L., Buenzli, P.R., Woodruff, M.A., Simpson, M.J., New computational tools and experiments reveal how geometry affects tissue growth in 3D printed scaffolds., Chemical Engineering Journal, 2023; 475: 145776. doi: 10.1016/j.cej.2023.145776.

Lassen, N.E., Andersen, T.L., Pløen G.G., Søe, K., Hauge, E.M., Harving, S., Eschen, G.E.T., Delaisse, J-M., Coupling of Bone Resorption and Formation in Real Time: New Knowledge Gained From Human Haversian BMUs., Journal of Bone and Mineral Research, 2017; 32: 1395–1405. doi: 10.1002/jbmr.3091.

Robling, A., Stout, S., Morphology of the drifting osteon., Cells Tissues Organs, 1998; 164: 192–204. doi: 10.1159/000016659.

Schnitzler, C.M., Mesquita, J.M., Cortical porosity in children is determined by age-dependent osteonal morphology., Bone, 2013; 55: 476–486. doi: 10.1016/j.bone.2013.03.021.

Cooke, K.M., Mahoney, P., Miszkiewicz, J.J., Secondary osteon variants and remodeling in human bone., The Anatomical Record, 2022; 305: 1299–1315. doi: 10.1002/ar.24646.

Hegarty-cremer, S.G.D., Borggaard, X.G., Andreasen, C.M., van der Eerden, B.C.J., Simpson, M.J., Andersen, T.L., Buenzli, P.R., How osteons form: A quantitative hypothesis-testing analysis of cortical pore filling and wall asymmetry., Bone, 2024; 180: 116998. doi: 10.1016/j.bone.2023.116998.

Tee, S.Y., Bausch, A.R., Janmey, P.A., The mechanical cell., Current Biology, 2009; 19(17): R745 – R748. doi: 10.1016/j.cub.2009.06.034.

Marotti, G., Zallone, A.Z., Ledda, M., Number, Size and Arrangement of Osteoblasts in Osteons at Different Stages of Formation., Calcified Tissues, 1976; 21: 96–101. doi: 10.1007/BF02546434.

Werner, M., Blanquer, S.B.G., Haimi, S.P., Korus, G., Dunlop, J.W.C., Duda, G.N., Grijpma, D.W., Petersen, A., Surface Curvature Differentially Regulates Stem Cell Migration and Differentiation via Altered Attachment Morphology and Nuclear Deformation., Advanced science, 2017; 1600347. doi: 10.1002/advs.201600347.

Gudipaty, S.A., Lindbolm, J., loftus, P.D., Redd, M.J., Edes, K., Davey, C.F., Krishnegowda, V., Rosenblatt, J., Mechanical stretch triggers rapid epithelial cell division through piezol., Nature, 2017; 118–121. doi: 10.1038/nature21407.

Ravasio, A., Cheddadi, I., Chen, T., Pereira, T., Ong, H.T., Bertocchi, C., Brugues, A., Jacinto, A., Kabla, A.J., Toyama, Y., Trepat, X., Gov, N., Neves de Almeida, L., Ladoux, B., Gap geometry dictates epithelial closure efficiency., Nature communications, 2015; 6, 7683. doi: 10.1038/ncomms8683.

Kuba, S., Simpson, M. J., and Buenzli, P. R. (2026). Stochastic model of curvature-controlled tissue growth incorporating mechanical cell interactions. GitHub repository. https://github.com/Shahak-Kuba/2026_Stochastic_Tissue_Growth

